# Heterogeneity and developmental dynamics of LYVE-1 perivascular macrophages distribution in the mouse brain

**DOI:** 10.1101/2021.10.08.463501

**Authors:** Marie Karam, Guy Malkinson, Isabelle Brunet

## Abstract

Brain perivascular macrophages (PVMs) belong to border-associated macrophages. PVMs are situated along blood vessels in the Virchow-Robin space and are thus found at a unique anatomical position between the endothelium and the parenchyma. Owing to their location and phagocytic capabilities, PVMs are regarded as important components that regulate various aspects of brain physiology in health and pathophysiological states. Here, used LYVE-1 to identify PVMs in the mouse brain. We used brain-tissue sections and cleared whole-brains to learn how they are distributed within the brain and across different developmental postnatal stages. We find that LYVE-1^+^ PVMs associate with the vasculature in a brain-region-dependent manner, where the hippocampus shows the highest density of LYVE-1^+^ PVMs. We show that their postnatal distribution is developmentally dynamic and peaks at P10-P20 depending on the brain region. We further demonstrate that their density is reduced in the APP/PS1 mouse model of Alzheimer’s Disease. In conclusion, our results show an unexpected heterogeneity and dynamics of LYVE-1^+^ PVMs, and support an important role for this population of PVMs during development and in regulating brain functions in steady-state and disease conditions.

## Introduction

Continuous brain activity is crucial in order to support body demands and needs. Such extensive activity requires constant and considerable supply of oxygen and nutrients, a task which is carried out by the brain’s vascular network. However, it also generates an extensive amount of metabolites and by-products that must be cleared away from the brain tissue in order to ensure a stable physiological environment^1^. Exchange of molecules across the vascular membrane between the vessel lumen and the parenchyma is tightly regulated because of the blood brain barrier^2^, thus a critical limitation in terms of waste-clearance is imposed. Currently, several mechanisms account for waste-clearance from the brain. The first are the meningeal lymphatic vessels (mLVs), which are part of the endothelial lymphatic system^3,4^. Waste in the brain itself is cleared by a different system, the glymphatic system, which is a paravascular system responsible for the exchange of cerebrospinal fluid (CSF) into and out of the brain around the vessels, in the Virchow-Robin space^5^. An additional clearance route along arterial perivascular spaces has also been demonstrated^6^. Nonetheless, it remains largely unknown how physiological homeostasis is maintained both globally and locally in the brain^1, 7^.

Brain perivascular macrophages (PVMs) are a subset of border-associated-macrophages (BAMs, also known as CNS-associated-macrophages, CAMs) which also comprise the meningeal macrophages (MMs) and choroid-plexus macrophages (CPMs) ^8, 9^. PVMs are found in the Virchow-Robin space around the vessel wall, at a strategical anatomical position between the endothelium and the parenchyma, and are thus regarded as critical in linking peripheral blood-borne signals and the CNS, notably in regulating immune-surveillance and fluid homeostasis^6, 9, 10,11^.

The embryonic origins of PVMs are documented ^8, 12^, but surprisingly little is known on how PVMs are distributed in the brain in postnatal steady-state conditions ^9^ and whether their distribution undergoes spatio-temporal changes until adulthood and under pathophysiological conditions. At the molecular level, PVMs express a set of well-defined markers, notably CX3CR1, CD206 (also known as MRC1) and lymphatic vessel endothelial hyaluronan receptor 1 (LYVE-1)^13^. The specificity of LYVE-1 as a PVM marker was recently demonstrated^14^ and the *lyve-1* locus was used to distinguish between a subset of PVMs and CX3CR1^+^ microglia^15^. Although their exact role remains to be elucidated, LYVE-1^+^ PVMs display phagocytic capabilities and have been shown to be involved in hypertension^13,16^, neuroinflammation^14, 15^ and stroke^17^.

Currently, characterization of BAMs in general, and PVMs in particular, relies on enrichment by dissociation, followed by detailed molecular analyses. Although this approach can reach single-cell sensitivity and is highly informative, it lacks the spatial dimension since the location of the cells in their native tissues is not preserved. Moreover, the tissue and organs often need to be pooled in order to increase the quality and quantity of molecules in the sample. Molecular analyses of smaller regions within the organ or the tissue are even more challenging for the same reason.

Here we study for the first time the distribution of LYVE-1 PVMs at a brain-wide scale by visualizing them in unprocessed tissue. We used immuno-labeling combined with iDisco tissue clearing^18^ of entire mouse brains to visualize the vasculature that is associated with LYVE-1^+^ PVMs. We examined different developmental postnatal periods from birth until adulthood, and compared the density of these PVMs in different brain regions. Our results demonstrate a heterogeneous distribution of LYVE-1+ PVMs in the brain, and we furthermore find that as the brain ages, the coverage fraction of the brain vasculature by these PVMs changes as well.

## Materials and Methods

### Animals

Male and female *C57Bl/6Nj* mice were purchased from Janvier Laboratories, and APP/PS1 mice (B6;C3-Tg(APPswe,PSEN1dE9)85Dbo/Mmjax) were also purchased from Jackson Laboratories. No selection criteria were used for selecting the animals. The mice were housed in temperature and humidity controlled rooms, maintained on a 12h/12h light/dark. All strains were kept in identical housing conditions in pathogen-free facility and were handled in compliance with regulations of the ethical rules of the French agency for animal experimentation. Experiments were approved by the French Ministry for Research and Higher Education’s institutional review board (Apafis number: 21814). Mice were euthanized at various ages ranging from 0-100 weeks of age. The experiments were performed in accordance with the ARRIVE guidelines. Sample size was chosen in accordance with similar, previously published experiment.

### Perfusion and tissue processing

Mice were euthanized with an intraperitoneal (i.p.) injection of 400mg/kg of ketamine (Imalgene) and 20mg/kg of Xylazine (Rompun) and then intracardiaclly perfused with 0.1M of PBS for 5 min until exsanguination followed by 4% paraformaldehyde (PFA) for fixation, and the brains were immediately extracted. After extraction, the brain was immersed in a 4% PFA solution, overnight for 24h at 4°C. The brains were then embedded in agarose and 100-150 μm brain sections were made using a vibration microtome (HM 650 V).

### Brain Immunolabeling

Brain sections were incubated with PBS containing 1% Triton-X-100 for 1h at room temperature (RT), followed by 3 washes (15min each) with PBS containing 10% tris-HCL PH 7,4, 3% NaCl 5M and 1% Triton-X-100 (TNT). The sections were then blocked with a blocking solution (TNBT) containing 10% Tris-HCL PH 7,4, 3% NaCl 5M, 1% Triton-X-100, 0,5% Perking blocking reagent all diluted in H_2_O milliQ QSP. This was then followed by an overnight incubation at 4°C with appropriately diluted primary antibodies: anti-CD31 (R&D Systems, clone AF3628, 1:250); anti-Lyve-1(ReliaTech GmbH, clone 103-PA50, 1:300), anti-Lyve-1 (eBioscience, ALY7, 1:100), anti-CD206 (Biorad MCA2235T, clone MR5D3, 1:300), anti-aquaporin 4 (Sigma, A5971, 1:500), anti-ephb4 (R&D Systems, AF446-SP, 1:100), anti-SMA (Sigma, C6198-2ML, 1 :250). Whole mounts and sections were then washed 3 times for 15 min at RT with TNT, followed by incubation with Alexa-fluor 488/555/647 Donkey/ anti rabbit/goat/rat IgG antibodies (Invitrogen, 1:250) for 4h at RT in TNBT. After 5 min in 1:2000 DAPI reagent, sections were washed 3 times with TNT and mounted with a fluorescence mounting medium (Dako, S3023) under coverslips.

### Samples staining and iDISCO+ clearing

Entire mice brains were cleared using the iDisco clearing protocol^18^. For the immunolabeling step, the brains were incubated for 5-7 days at 37°C with rotation, with diluted primary antibodies: anti-CD31 (R&D Systems, clone AF3628, 1:400), anti-Lyve-1(ReliaTech GmbH, clone 103-PA50, 1:330), anti-Ephb4 (R&D Systems, AF446-SP, 1:100). Followed by 4 days incubation at 37°C with rotation, with diluted secondary antibodies: Alexa-fluor 555/647 Donkey anti Rabbit/Goat IgG antibodies (Invitrogen, 1:400).

### Brain slices imaging

Images of brain sections were taken using Zeiss Axiozoom apotome, and Zeiss LSM 980 confocal microscope.

### Light sheet imaging

Brains were imaged either on the Ultramicroscope II (LaVision Bio Tec) or Ultramicroscope Blaze (Miltenyi Biotec) light-sheet microscope with a 1.0X objective, and on a light-sheet microscope with a 0.63X objective.

### Image processing and quantification method

For figure 1I, using imagej software, a thresholding step was made for both LYVE-1 and CD206 channel, then both particles were analyzed and the number of count was extracted.

**Figure 1:**
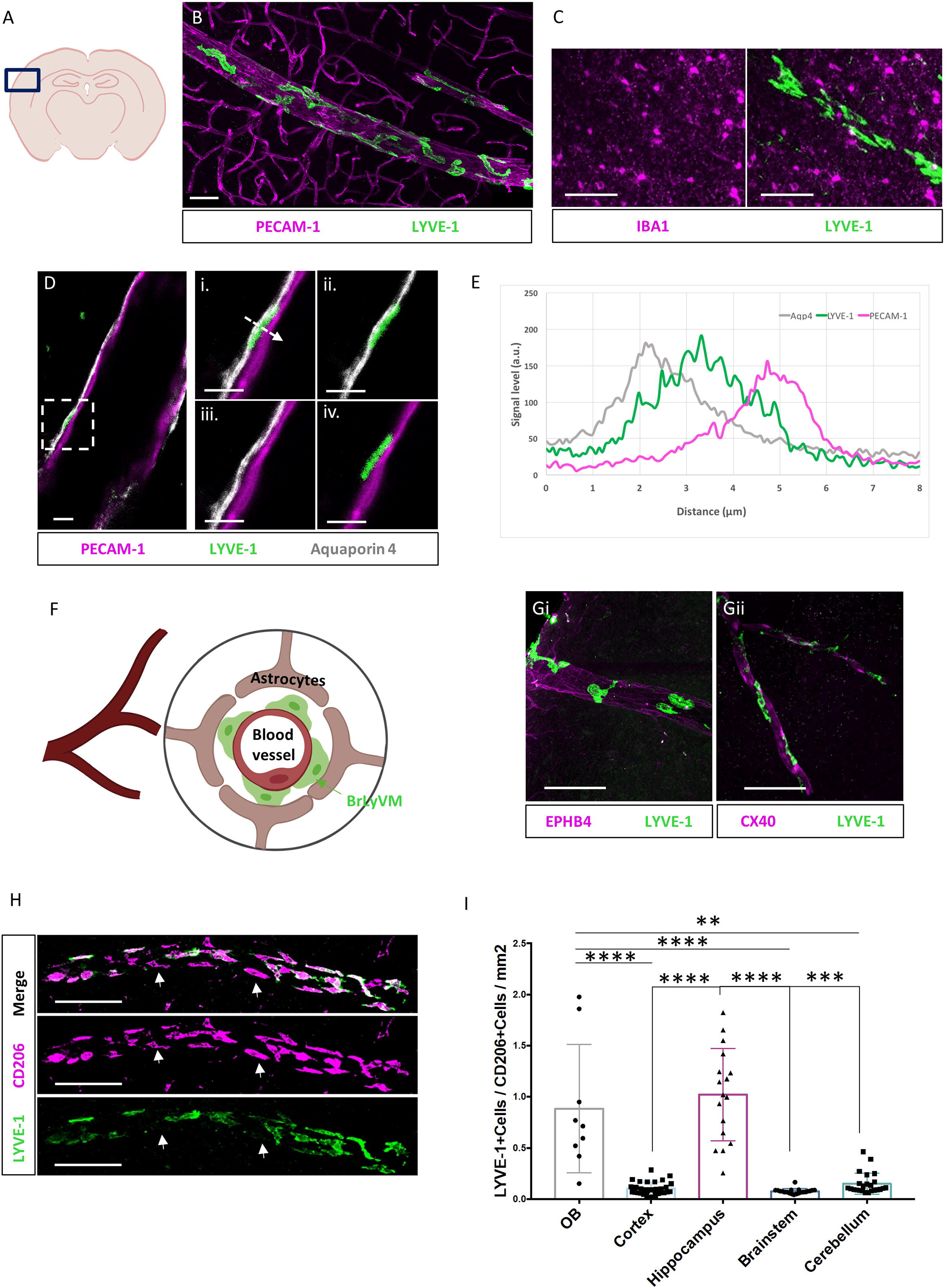
Characterization of the vascular environment of BrLyVMs. **(A)** Schematic representation of a brain slice showing the region that was used for generating the images. **(B)** Immuno-labeling of LYVE-1 (green) and PECAM-1 (magenta) reveals that BrLyVMs are not part of the endothelium. Scale bar 50µm. **(C)** Immuno-labeling of LYVE-1 (green) and microglia (IBA1, magenta) demonstrates that BrLyVMs and microglia are morphologically distinct. Scale bar 50µm. **(D)** BrLyVMs (LYVE-1, green) are positioned between the endothelium (PECAM-1, magenta) and the glia limitans (Aquaporin 4, white). **(Di-Div)** Higher magnification of the box highlighted in (**D**). Scale bar 10µm. (**E**) Graphical representation of the three different markers along the dashed line shown in (**Di**). **(F)** Schematic representation of the localization of BrLyVMs in the Virchow-Robin space. Astrocytes, BrLyVMs and a blood vessel are annotated. **(G)** BrLyVMs encircle veins (EPHB4, magenta, **Gi**) and arteries (CX40, magenta, **Gii**). Scale bar 100µm. **(H)** Co-immunolabeling for LYVE-1 (green) and CD206 (magenta). Scale bar 100µm. Arrows point to cells that express CD206 but are negative for LYVE-1. **(I)** Quantification of BrLyVMs as a fraction of CD206^+^ PVMs in different brain regions. (mean ± SD; n = 3 mice each group; *p<0.05 Kruskal-Wallis with Dunn’s post-hoc test).

For figure 2, 4 and 5, Imaris software was used to process the brain by extracting in 3D its different regions (OB, Cortex, Hippocampus, Brainstem and Cerebellum) using contour surface, that allowed to manually select the brain region contours on 2D slices while precisely and specifically removing the pial surface. This was followed by a mask step for the created surface.

**Figure 2:**
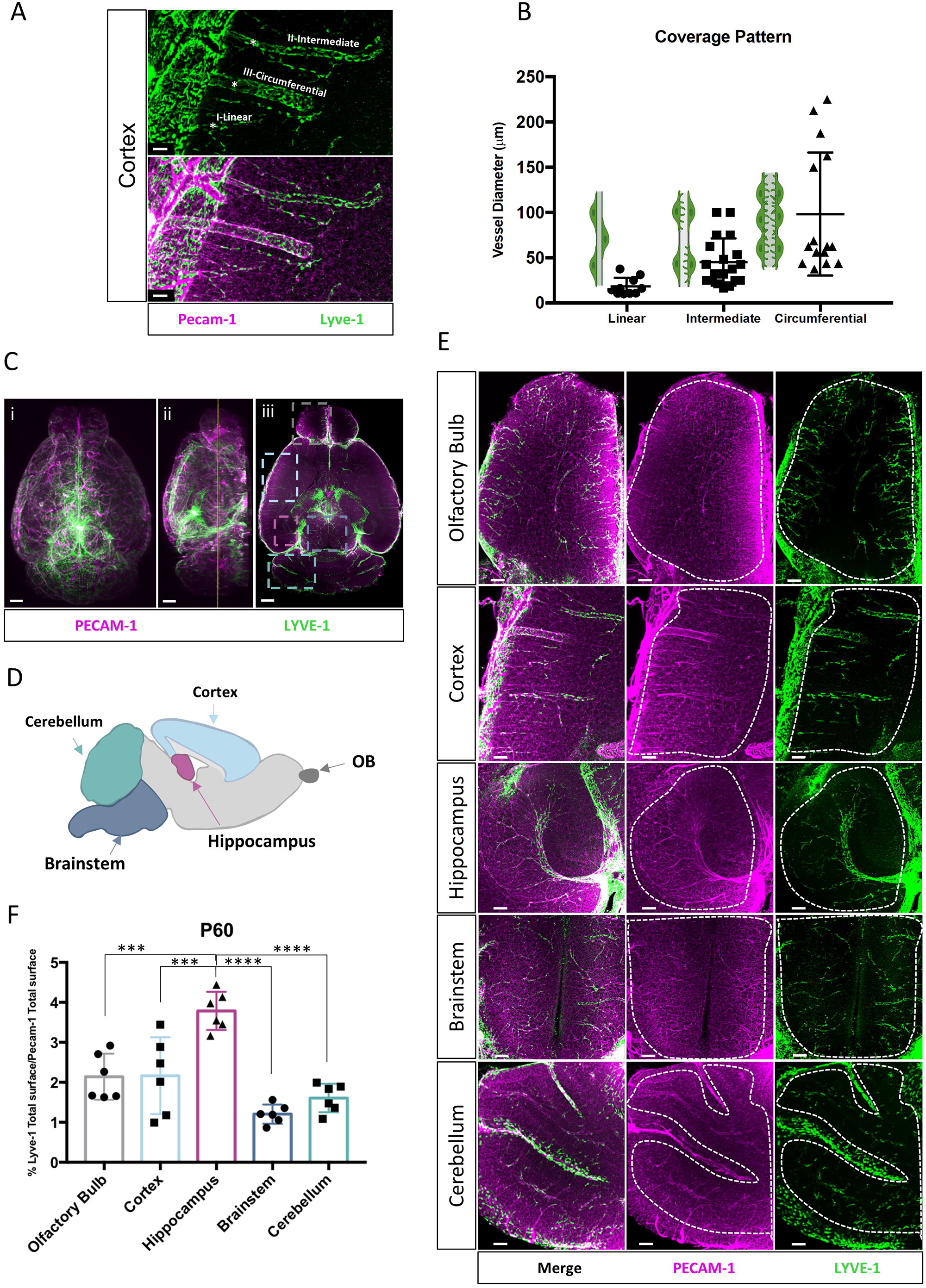
The heterogeneity of BrLyVMs distribution and density in the adult mouse brain revealed by tissue clearing. (**A**) Three dimensional (3D) rendering of a 500µm-thick optical slice of the cortex immunolabeled for PECAM-1 (magenta) and LYVE-1 (green) after iDisco tissue clearing. Three different coverage patterns around the endothelium are observed (denoted as I, II and III on the image). (**B**) Quantification of the three patterns as a function of vessel diameter. Shown in green are schematic drawings that represent each pattern. n=3 (**C**) 3D renderings of dorsal (**i**), lateral (**ii**) and a single-optical slice (**iii**) views of LYVE-1 (green) and PECAM-1 (magenta) wholemount immunolabeling. Scale bar 700µm. (**D**) Schematic annotated representation of five brain regions shown in color-coded dashed boxes in (**Ciii**). (**E**) Higher magnification of the boxes highlighted in (**Ciii**) representing 500µm-thick optical slice. Dashed lines encircle the regions that were used for the analysis. Scale bar 200µm. (**F**) Quantification of LYVE-1 surface area as a fraction of total PECAM-1 surface area in each region (mean ± SD; n = 6 mice each group; *p<0.05 one-way ANOVA with Bonferroni post-hoc test).

**Figure 3:**
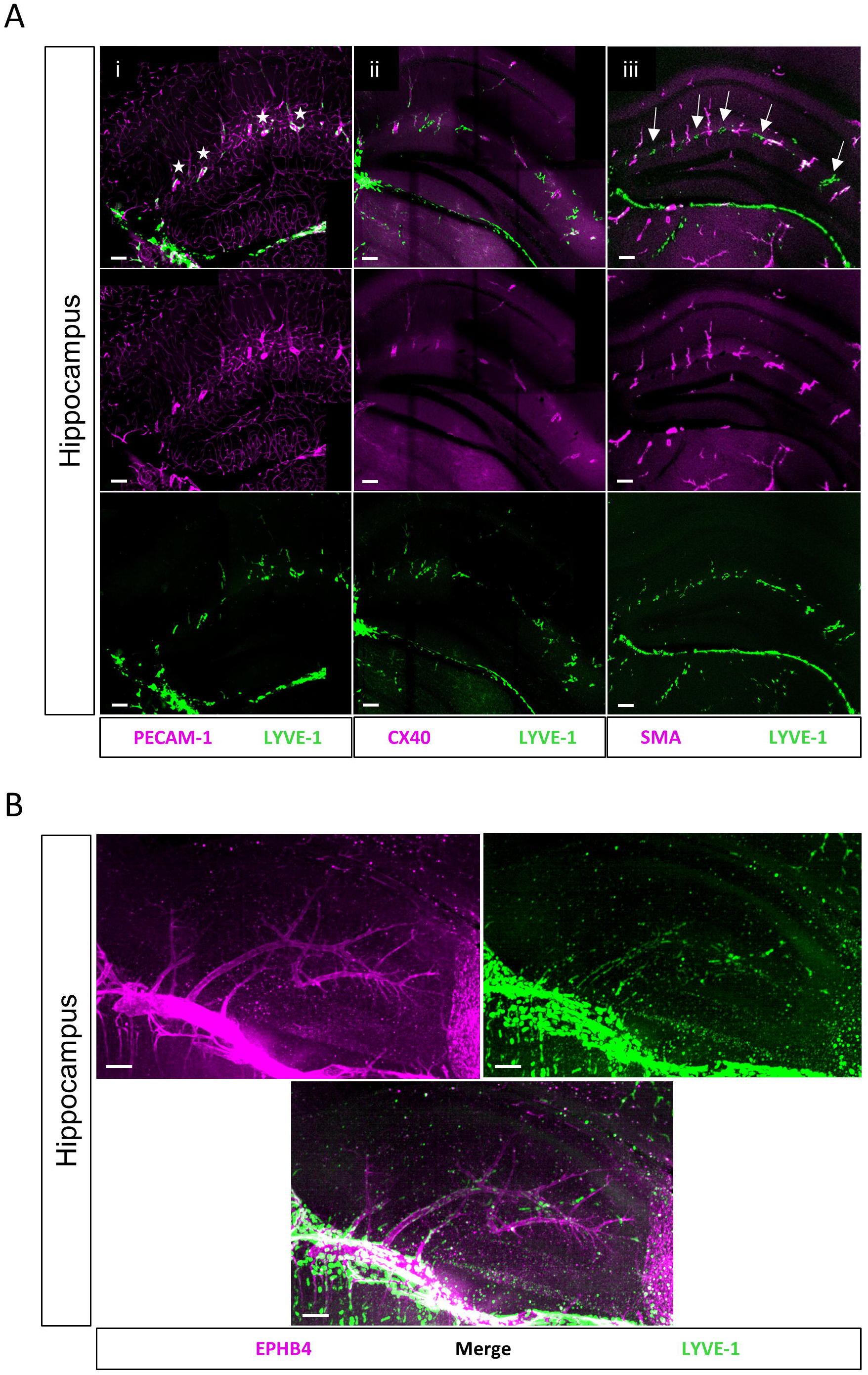
Hippocampal BrLyVMs encircle main arteries and veins. (**A**) Hippocampal region of brain slices co-immunolabeled for LYVE-1 (green) and PECAM-1 immunolabeling (magenta, **Ai**), CX40-GFP (magenta, **Aii**) or smooth-muscle-actin immunolabeling (SMA, magenta, **Aiii**). BrLyVMs encircle large-caliber hippocampal vessels (white stars, **Ai** top row). Arteries (**Aii, Aiii**) are encircled by BrLyVMs. White arrows (**Aiii**, top row) denote LYVE-1 around veins that are detected as SMA-negative vessels in between SMA-positive vessels. Scale bar 200µm. (**B**) Immunolabeling of whole-mount cleared brain tissue for LYVE-1 (green) and EPHB4 (magenta) reveals large-caliber veins that are encircled by BrLyVMs (scale bar = 200µm).

**Figure 4:**
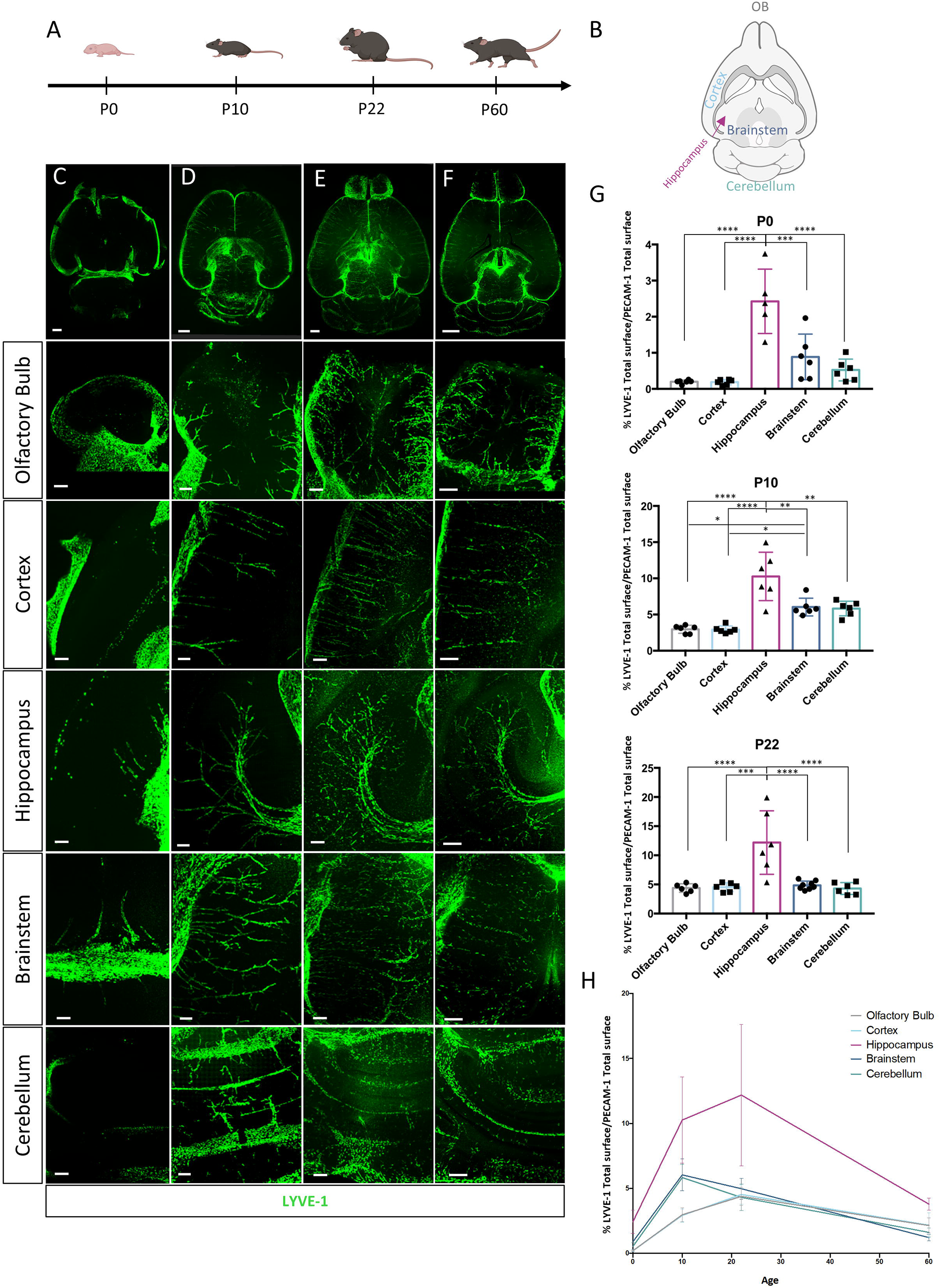
BrLyVMs dynamics at different developmental stages revealed by tissue clearing. (**A**) Schematic representation of the different developmental stages examined. (**B**) Schematic representation of the different regions that were used for imaging and for quantification. (**C-F**) Representative 200µm optical-slice dorsal views of LYVE-1 (green) wholemount immunolabeling of the olfactory bulb, cortex, hippocampus, brainstem and cerebellum at the different stages. Columns **C, D, E, F** correspond to P0 (scale bar 100µm), P10 (scale bar 200µm), P22 (scale bar = 200µm) and P60 (scale bar = 200µm), respectively. (**G**) Quantification of LYVE-1 surface area as a fraction of total PECAM-1 surface area in each of the different brain regions at the different developmental stages (mean ± SD; n = 6 mice each group; *p<0.05 one-way ANOVA with Bonferroni post-hoc test). (**H**) Representative graph of the BrLyVMs dynamics in each quantified brain region at the different time points. The peak density in the brain stem and in the cerebellum is seen at P10, while that of olfactory bulb, cortex and hippocampus is seen at P22.

**Figure 5:**
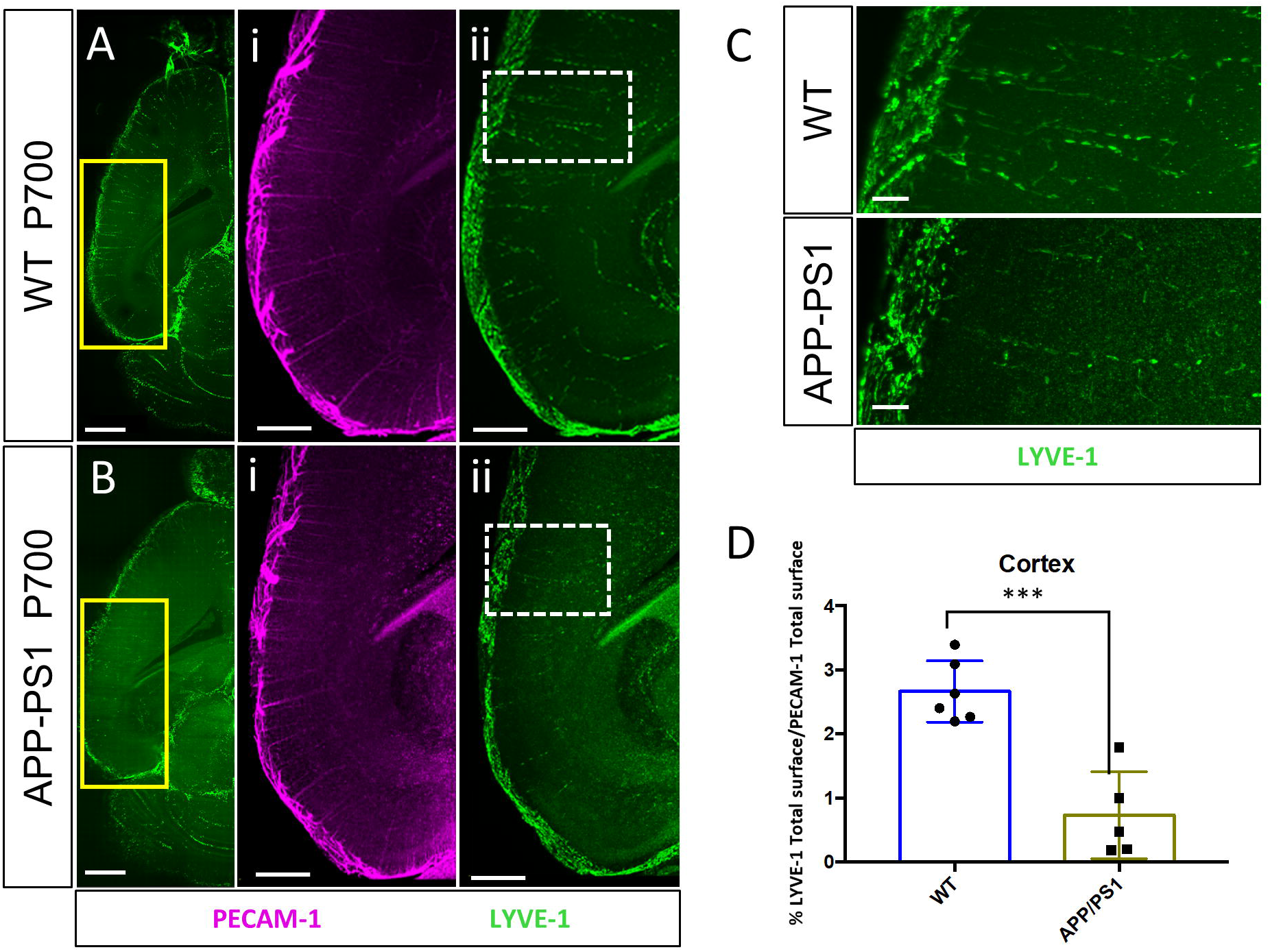
BrLyVMs density in pathophysiological conditions revealed by tissue clearing. (**A, B**) Representative 200µm optical slice of LYVE-1 (green) and PECAM-1 (magenta) wholemount immunolabeling of WT siblings (**A**) and APP/PS1 (**B**) brains at P700, respectively (dorsal view). Scale bar 1000µm. (**Ai, Aii, Bi, Bii**)-higher magnification of the boxes highlighted in (**A, B**, scale bars = 500µm). (**C**) Higher magnification of the boxes highlighted in Ai and Bi (scale bars = 100µm). (**D**) Quantification of LYVE-1 surface area as a fraction of total PECAM-1 surface area in the cortex of P700 WT siblings and APP/PS1 mice (mean ± SD; n = 6 mice for the WT group and n = 5 for the APP/PS1 group; *p<0.05 student-t test).

For the quantification process, the brain vessels and Lyve-1 PVMs were identified, segmented and measured (area and volume) using Surfaces. The ratio calculated to generate Lyve-1 PVMs region and age dependent density were:

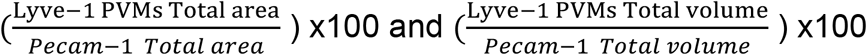

For Figure 5, analysis was performed without prior knowledge of group allocation.

### Statistical analysis

All statistics were conducted using using GraphPad Prism software. Throughout this article, data are represented as means with standard deviation. For figure 1, a non-parametric Kruskal-Wallis test was performed followed by a Dunn’s multiple comparisons test. For figure 2 and 4, a parametric one way Anova test was performed followed by Bonferroni post-hoc test for the mean comparisons between each group. For Figure 5, a parametric student t-test was used. All sample sizes and p values are indicated in the figure legends. In all instances, n represents the number of mice used.

## Results

To learn more about the immediate environment in which brain LYVE-1 PVMs (hereafter referred to as “BrLyVM”) are situated, we co-immunolabeled PECAM-1 to label the endothelium and LYVE-1 to label BrLyVMs. LYVE-1 signal did not co-localize with PECAM-1-labeled endothelium but rather enveloped it, confirming a perivascular localization of BrLyVM (Figure 1A-B, and Supplementary Video 1). While PVMs and parenchymal microglia are known to have common molecular markers^19^, BrLyVMs were elongated and morphologically distinct from microglia, which had a punctate-like staining pattern, as seen by co-immunolabeling for LYVE-1 and for the microglial marker Iba-1 (Figure 1C). To better understand where in the perivascular space BrLyVMs are situated, we stained for PECAM-1, LYVE-1 and Aquaporin 4, and found that BrLyVMs were situated between the endothelium and the glia limitans (Figure 1D). The signal level of the staining was also measured, confirming the perivascular localization of these BrLyVMs (Figure 1E-F).

The existing uncertainty in the literature regarding the distribution of PVMs around arteries and veins^9^ led us to next ask whether BrLyVMs are found around veins or arteries. We detected BrLyVMs around veins (Figure 1Gi) by co-labeling for LYVE-1 and for EPHB4, an established venous marker^20,21^. We used the transgenic mouse reporter line CONNEXIN-40-GFP^22^ (CX40-GFP) in which arterial endothelial cells express the fluorescent protein GFP. We co-immunolabeled CX40-GFP (GFP positive cells are displayed in magenta) brain slices for LYVE-1 and found, as previously reported, that BrLyVMs encircle arteries (Figure 1Gii). BAMs are known to express CD206^23^, also known as MRC1, but the relative abundance of these two markers in brain tissue has never been visualized *in situ*. We thus co-immunolabeled brain slices for these two markers (Figure 1H) and quantified their relative proportions in five different brain regions. We found that the ratio of BrLyVMs/CD206^+^ PVMs significantly varied between the different regions that we analyzed. In the olfactory bulb and in the hippocampus there was a significantly higher percentage of BrLyVMs/CD206^+^ PVMs than in the cortex, brainstem and cerebellum (Figure 1I).

Taken together, these results show that BrLyVMs are situated between the endothelial basement membrane and the glia limitans, which delimits the parnenchyma, and furthermore suggest that they possess unique molecular signatures and spatial distribution across brain regions.

PVMs are not abundant in the brain^8^, and furthermore we noticed that BrLyVMs abundance in the cortex was sparse and occasionally varied between brain regions (not shown). This raised the hypothesis that across different brain regions, BrLyVMs may be associated to different extents with the vasculature, which prompted us to investigate their heterogeneity at a global brain-wide scale. We used the iDisco tissue-clearing method that enables to detect and visualize large volumes of tissue at high resolution^18^. We verified the binding specificity of the LYVE-1 antibody with iDisco (Supplementary Figure 1A), and then co-immunolabeled entire brains with PECAM-1 and LYVE-1 (Figure 2). Inspection of immuno-labeled cleared cortical tissue revealed that BrLyVMs covered the endothelium (Figure 2A), in agreement with our results obtained on tissue slices (Figure 1). We further noticed that BrLyVMs encircled the cortical vasculature (hereafter, for the purpose of readability, we refer to any vessel that is encircled by BrLyVMs as “LYVessel”) in one of three coverage patterns (Figure 2A). Pattern I (“linear”) refers to vessels that were typically 10-40 µm microns in diameter and were characterized by a single line of BrLyVMs around their perimeter (Figure 2A, 2B). Pattern II (“intermediate”) vessels were of 20-100µm diameter, and contained more than one BrLyVM around their cross section (Figure 2A, 2B). In addition, class II vessels BrLyVMs displayed an irregular cellular shape. Pattern III (“circumferential”) vessel diameters ranged between 40-225µm, and showed a higher density of coverage and an irregular cellular shape (Figure 2A, 2B).

We were next curious to understand how BrLyVM are distributed in the brain and whether this distribution is homogenous in nature. Indeed, while examining the cleared immunolabeled brains we noticed that although the fluorescent signal was not ubiquitous as that of PECAM-1 (Figure 2C) across the brain, it was nevertheless detectable in specific regions in the brain (Figure 2C-D). We detected LYVE-1 staining in prominent brain structures: the cortex, hippocampus, olfactory bulb, cerebellum and brainstem, indicative of efficient antibody labeling and penetration across the brain. We visually identified gross-level differences in the labeling distribution (Figure 2E). To learn more about these differences and about if and how different brain regions differ in BrLyVM density, we proceeded by quantifying our datasets after removal of pia LYVE-1 signal (Supplementary Figure 2). We reasoned that in order to exclude anatomical intra-regional differences in vasculature density, it was necessary to normalize the density of BrLyVMs to that of the vasculature in each region. We thus calculated the fraction of the vasculature that is associated with BrLyVMs by normalizing the surface area of LYVE-1 to that of PECAM-1 in each of the examined regions (see Materials and Methods). We analyzed the olfactory bulb, cortex, hippocampus, brain stem and cerebellum, and found that BrLyVM density varied across these regions and was significantly higher in the hippocampus (Figure 2F).

Given that little is known about how BrLyVMs are distributed around the vasculature in the hippocampus, together with the relatively higher density of hippocampal BrLyVMs that we found (Figure 2), prompted us to examine this region in more detail. We noticed that hippocampal BrLyVMs were distributed in a stereotypical pattern (Supplementary Video 2). To learn more about this pattern, we first co-stained PECAM-1 and LYVE-1 and detected a preferential BrLyVMs association to large-caliber hippocampal vessels (Figure 3Ai). We then investigated if in the hippocampus, BrLyVMs are associated with arteries or veins. We co-labeled for LYVE-1 and for arteries, using either the CX40-GFP transgenic mouse line (Figure 3Aii) or by co-immunolabeling for smooth-muscle-actin (SMA, also known as acta2) to detect the arterial wall^24, 25^ (Figure 3Aiii). The results showed that in the hippocampus, BrLyVMs are associated with most if not all arteries. Arrows in Figure 3Aiii indicate LYVE-1 staining that alternates with arteries within the hippocampus. We reasoned that this staining could be BrLyVMs that are associated with veins, thus to see whether BrLyVMs are found around hippocampal veins, we analyzed cleared brain samples co-immunolabeled for LYVE-1 and for EPHB4. We found that and found that BrLyVMs are also present around large-caliber hippocampal veins (Figure 3B).

The heterogeneity that we observed in different brain regions (Figure 2) led us to hypothesize that BrLyVM are dynamic and spatially-regulated over time. The abundance of BAMs and of PVMs in different embryonic mouse stages was recently shown to be developmentally regulated^26^, and we hypothesized that BrLyVMs would display a dynamic organization from early postnatal stages into adulthood. In order to examine this hypothesis, we used whole-brain iDisco clearing of entire brains at P0, P10, P22 and P60 (Figure 4A-4F and Supplementary figure 3, Supplementary Figure 4 and Supplementary Figure 5A-D) and quantified BrLyVMs density as before (Figure 4G, 4H and Supplementary Figure 5E). We found that at P0, the majority of the LYVE-1 signal was found in the pia (Figure 4C and Supplementary Video 3). Although LYVessels were generally absent at this stage, we could nevertheless identify some LYVessels, notably in the brain stem and in the hippocampus (Figure 4C Brainstem, Hippocampus). At P10 and P22 (Figures 4D and 4E respectively) we saw a significant increase in LYVE-1 signal that was concomitantly associated with the vasculature, in the examined brain regions (Figure 4G and Supplementary Figures 3, 4 and 5, and Supplementary Video 4). The quantification revealed that while the density of BrLyVMs in the brainstem and the cerebellum peaks at around P10, the same density reaches its peak at around P22 in the other examined regions, namely the olfactory bulb, cortex and hippocampus (Figure 4G-H and Supplementary Figure 5E). Interestingly, a small but significant decrease in the density was noted in all regions by P60 (Figures 2F, 4G-H and Supplementary Figure 5E). The results that we observed during the different developmental stages (Figure 4) suggested that BrLyVM density and number are not constant but are rather dynamically regulated in response to the physiological condition and age of the animal. Interestingly, among the areas investigated, two distinct temporal dynamics were identified, and the hippocampus showed a BrLyVM density that is significantly higher than the one found in all other brain areas over time.

PVMs in the cortex are known to have a role in regulation of vascular amyloid-beta plaques deposition^27^. We thus asked whether we could detect alterations in cortical BrLyVMs density in a mouse model of Alzheimer’s Disease (AD), namely the APP/PS1 transgenic line in which amyloid beta plaques develop in the cortex (Supplementary Figure 6). We performed PECAM-1 and LYVE-1 immunolabeling on entire brains of 700 days-old mice followed by tissue clearing (Figure 5A-C). We quantified BrLyVM density after normalizing to the density of the vasculature, and found that it was significantly reduced in APP/PS1 mutants compared to their wild type siblings (Figure 5D), indicating that the population of BrLyVMs is altered in the cortex these mice.

## Discussion

Here we studied the postnatal distribution of BrLyVMs across the mouse brain by examining different stages from birth until adulthood. We used a brain-wide visualization approach that enabled us to efficiently detect the brain endothelium that is associated with this population of PVMs in intact brains. This methodology proved to be highly reliable and recapitulated hallmark properties of PVMs, establishing this approach as bona-fide for the study of BrLyVMs. Our results demonstrate for the first time that BrLyVMs occupy different proportions of the vasculature in different brain regions. Overall, a small percentage of brain vasculature is associated with them, although we noticed that some vessels are encircled for considerable lengths, reaching a few millimeters in certain cases (not shown). We also find that LYVessels are encircled by BrLyVMs in one of three patterns, depending on the diameter of the LYVessel. We show that there is a considerable difference in BrLyVM density during the first postnatal weeks in a brain-region-dependent manner. BrLyVMs massively populate the brain between P0 to P10, peaking at around P10 or P20, depending on the brain region. Finally, our results demonstrate a transition in the density and morphology of BrLyVMs in a pathophysiological mouse model of AD, in the cortex where amyloid beta plaques develop. Whether plaque deposition is altered by a diminution of BrLyVMs remain to be deciphered.

The anatomical data we obtained support the view that BrLyVMs are not abundant in the brain^8^, and that they are associated with a small fraction of the vasculature, which suggests a low density of LYVessels in the brain. This raises the possibility that LYVessels have a specific signature that renders them different from neighboring non-LYVessels. This signature could be molecular or functional, or both, and remains to be elucidated, but in any case, our observations are in agreement with the known diversity of BAMs^28^. Thus it is also likely that different sub-classes of PVMs, as identified by the different markers, associate with different vessels and that their ensemble yields a more significant coverage of the vasculature. The variability that we see in the coverage density between different brain regions is intriguing and is also in-line with previous reports showing that brain endothelium exhibits considerable molecular and functional heterogeneity between different brain regions^29,30^ and even within-region differences^25, 31^. In this regard, it is interesting to note that LYVessels can be either arterial or venous. This suggests that arterial and venous BrLyVMs may be molecularly distinct, in line with the molecular heterogeneity of arterial and venous endothelial cells ^32^.

Our samples also enabled us to detect for the first time different coverage patterns in which BrLyVMs encircle the endothelial wall. The detection of these patterns was possible because the structural integrity of the vascular tissue remained intact. The patterns we observed suggest that there are mechanisms that link certain properties of the vessel to the coverage pattern. Indeed, why a certain pattern is associated with a certain vessel remains to be explored more in detail, but could be linked to the diameter of the vessel, to its arterial or venous identity, or to other unknown properties. Nonetheless, this observation strongly supports an important role for PVMs in contributing to vascular aspects of brain physiology.

We find that at P0, most of the LYVE-1 signal is associated with the pia, although a small number of LYVessels can be detected in specific brain regions, namely the brain stem and the hippocampus. Our results are in agreement with low PVM numbers that are found in E18.5 brains^26^. Between P0 and P10 there is a significant increase in LYVE-1 signal inside the brain, where the signal is detected in BrLyVMs. This suggests that between these two time points there is a massive invasion of BrLyVMs into the brain tissue, in a manner that is similar to the way that microglia colonize the brain until the second postnatal week in mice^33, 34^. From a developmental point of view, our results substantiate the reported cellular plasticity of BAMs during the first few weeks of postnatal development^28^.

So far, BAMs have been characterized mostly in mice, but also in zebrafish and in non-human primates (NHP). Zebrafish BAMs comprise mainly MMs^35,36^, and to a lesser extent PVMs and CPMs^37^. Interestingly, zebrafish BAMs are CD206^+^/LYVE-1^+^, and in NHP, CD206^+^ PVMs respond to viral infections^38, 39^. An across-species molecular and anatomical comparison of BAMs in general, and of BrLyVMs specifically, is expected to yield important insights about the evolution and ontogeny of these cells, and about their functional roles.

In conclusion, we provide here evidence to show that BrLyVMs are unevenly distributed across the brain, and that their distribution changes with age. This population of PVMs is plastic and dynamic overtime. Taken together with the existing evidence about the role of PVMs in general and BrLyVMs in particular, we propose that they play a significant role in brain physiology through currently unidentified interactions with the brain vascular system.

## Acknowledgements

The authors would like to thank members of the Brunet lab for technical assistance, Sara Makhoul for assistance with preparing the schematic illustrations and the Orion imaging facility, CIRB, for their support with the imaging presented in this article.

## Authors’ contributions

MK, GM and IB contributed to the conceptualization and experimental design.

MK performed experiments and analyzed the data.

All authors interpreted the analyzed data.

GM drafted the manuscript.

All authors edited, revised and approved the final manuscript.

## Supplementary material

There is supplementary material associated with this article

## Supplementary Figures

**Supplementary Figure 1**. A. Three dimensional (3D) rendering of a 500µm-thick optical slice of the cortex immunolabeled for i. LYVE-1 antibody (green). ii. LYVE-1 isotype control. iii. LYVE-1 secondary antibody. (scale bar = 500µm). Image intensity was linearly enhanced post-acquisition in (Aii, Aiii) for visualization purposes.

**Supplementary Figure 2**. (A) Manual drawing, using ‘‘Contour’’ on Imaris software, of the five quantified brain regions (i. Cortex, ii. Hippocampus, iii. Cerebellum, iv. Olfactory bulb, v. Brainstem) with the removal of the pial surface. (B) Masking of the contoured surface of i. Cortex, ii. Hippocampus, iii. Cerebellum, iv. Olfactory bulb, v. Brainstem. (C) The five quantified brain regions without the pial surface.

**Supplementary Figure 3**. Representative 200µm optical-slice dorsal views of LYVE-1 (green) and PECAM-1 (magenta) wholemount immunolabeling of the olfactory bulb, and the cortex at different stages. A. P0 (scale bar 100µm), B. P10 (scale bar 200µm), C. P22 (scale bar 200µm) and D. P60 (scale bar 200µm).

**Supplementary Figure 4**. Representative 200µm optical-slice dorsal views of LYVE-1 (green) and PECAM-1 (magenta) wholemount immunolabeling of the Hippocampus, and the Brainstem at different stages. A. P0 (scale bar 100µm), B. P10 (scale bar 200µm), C. P22 (scale bar 200µm) and D. P60 (scale bar 200µm).

**Supplementary Figure 5**. Representative 200µm optical-slice dorsal views of LYVE-1 (green) and PECAM-1 (magenta) wholemount immunolabeling of the cerebellum at the different stages. A. P0 (scale bar 100µm), B. P10 (scale bar 200µm), C. P22 (scale bar 200µm) and D. P60 (scale bar 200µm). E. Quantification of LYVE-1 surface area as a fraction of total PECAM-1 surface area at different developmental stages in the different brain regions (mean ± SD; n = 6 mice each group; *p<0.05 one-way ANOVA with Bonferroni post-hoc test).

**Supplementary Figure 6**. A. Representative figures of Thioflavin S staining in the cortex of a P700 WT and an APP/PS1 mice siblings (scale bar=100µm). White arrow points to an amyloid beta in Aii. B. the quantification of the average number of amyloid beta plaques in the cortex /mm2. (n=2 WT mice and n=3 APP/PS1 mice were used).

## Supplementary Videos

**Supplementary Video 1**. Three-dimensional animation of a PECAM-1 (magenta) immunolabeled vessel encircled by BrLyVMs detected with immuno-labeling against LYVE-1 (green).

**Supplementary Video 2**. Three-dimensional animation of a Hippocampus at P60 brain immunolabeled against LYVE-1 (green).

**Supplementary Video 3**. Three-dimensional animation of an entire P0 brain immunolabeled against LYVE-1 (green). The video zooms on a few BrLyVMs in the brainstem.

**Supplementary Video 4**. Three-dimensional animation of an entire P10 brain immunolabeled against LYVE-1 (green).

