## Supplementary figures and images for "Heterogeneity and developmental dynamics of LYVE-1 perivascular macrophages distribution in the mouse brain"

### Supplementary Figure 1

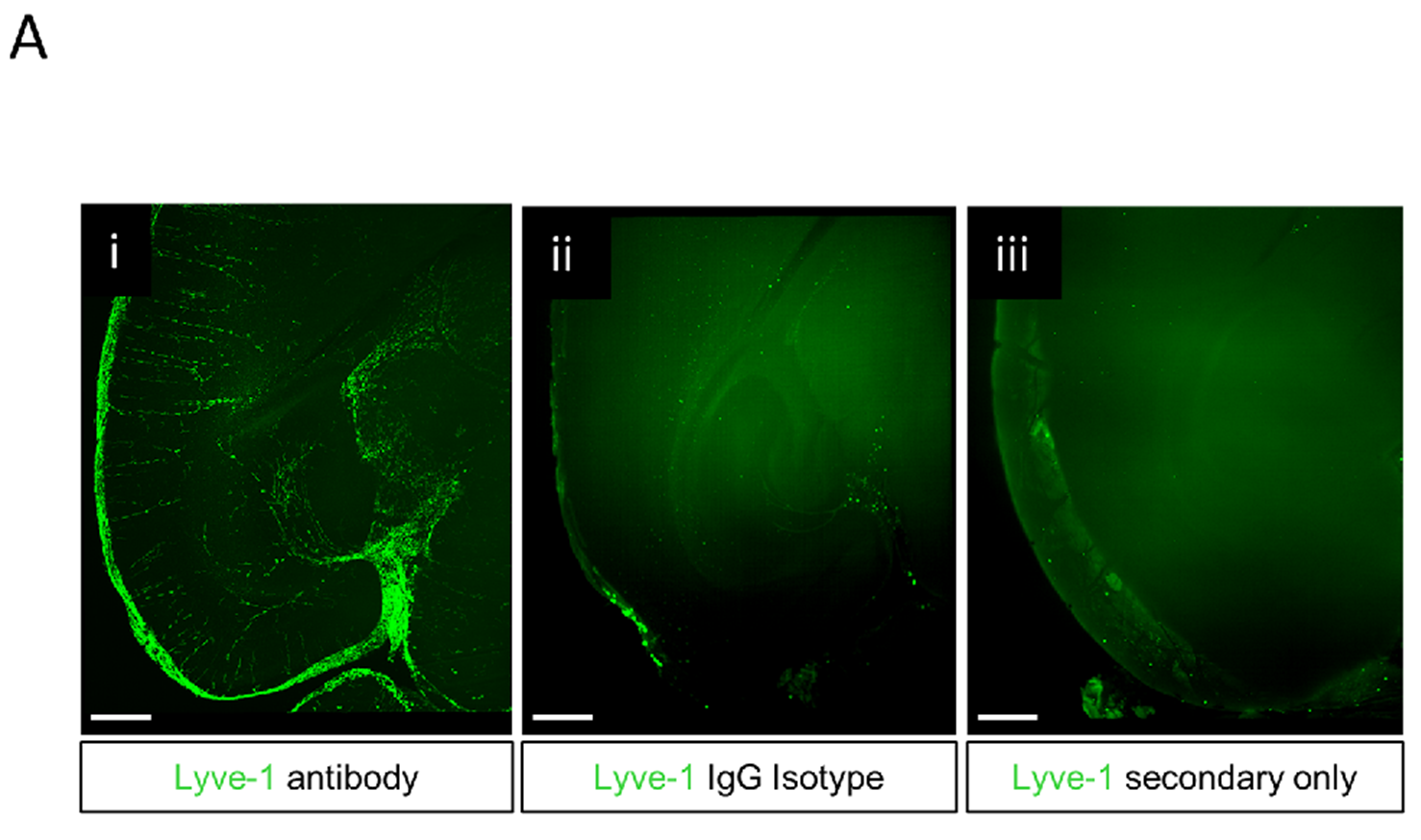

### Supplementary Figure 2

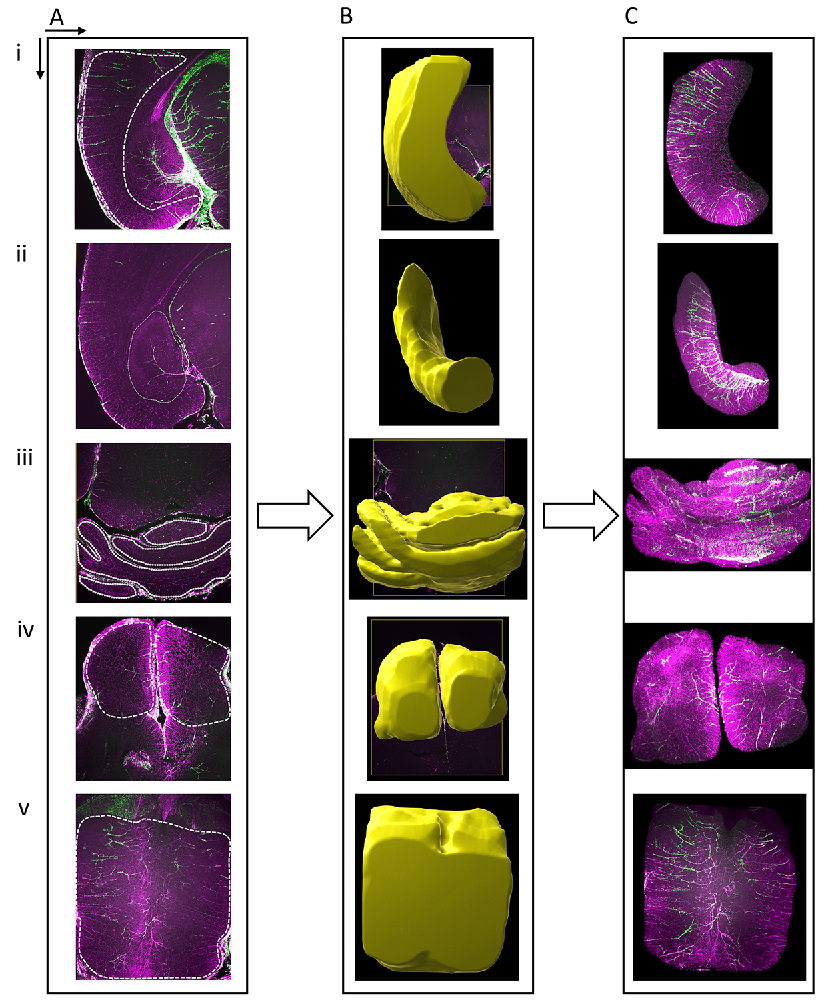

### Supplementary Figure 3

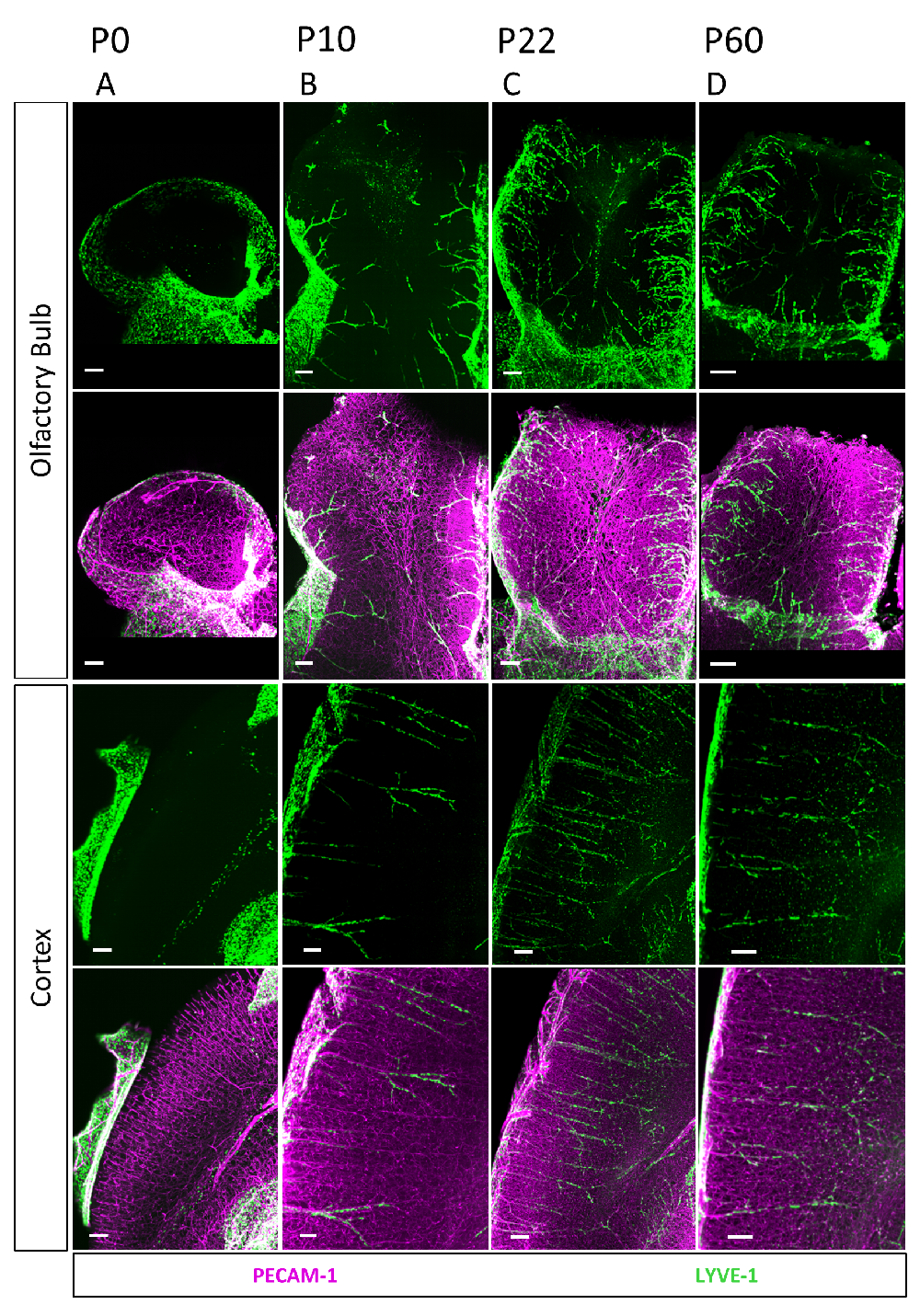

### Supplementary Figure 4

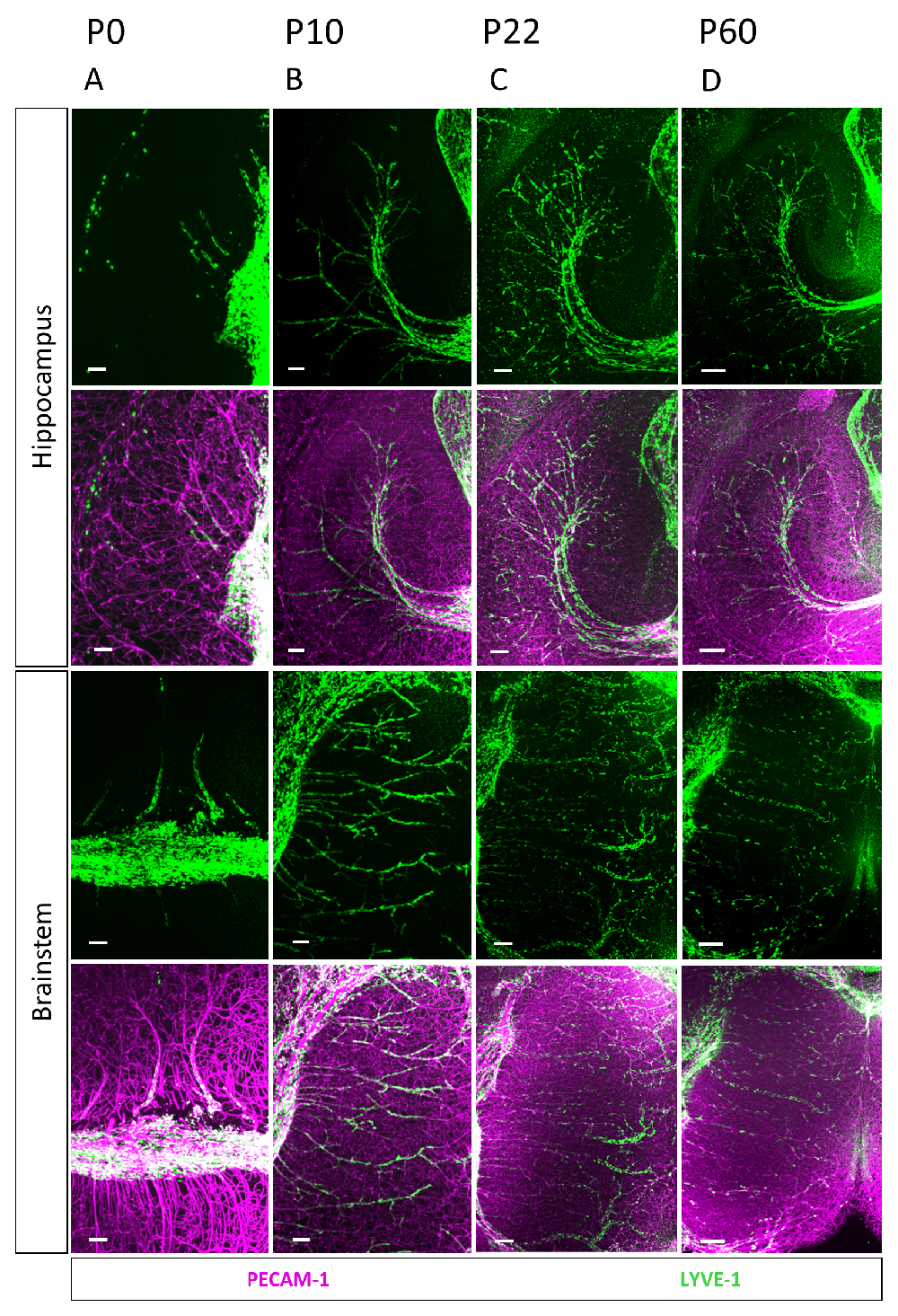

### Supplementary Figure 5

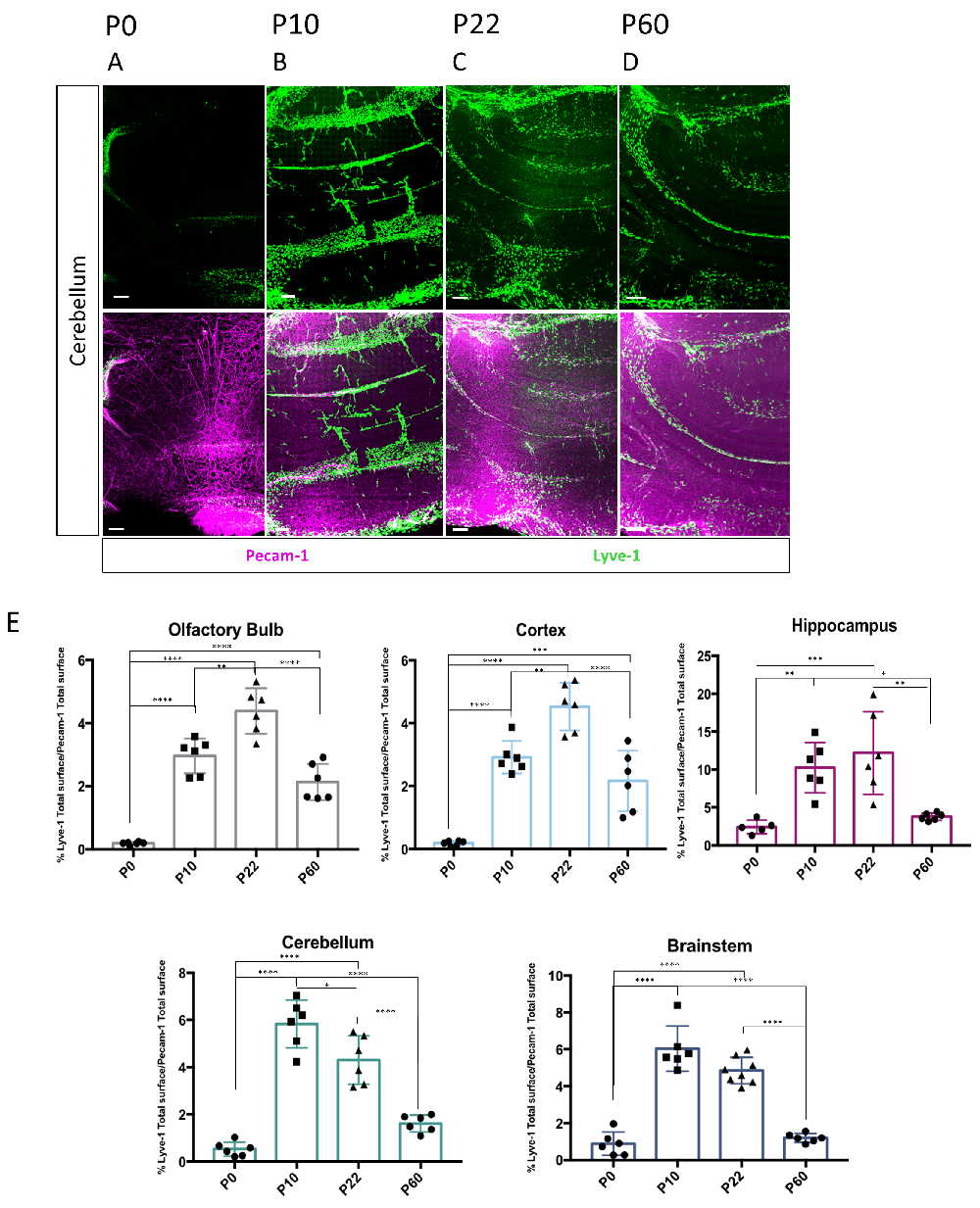

### Supplementary Figure 6

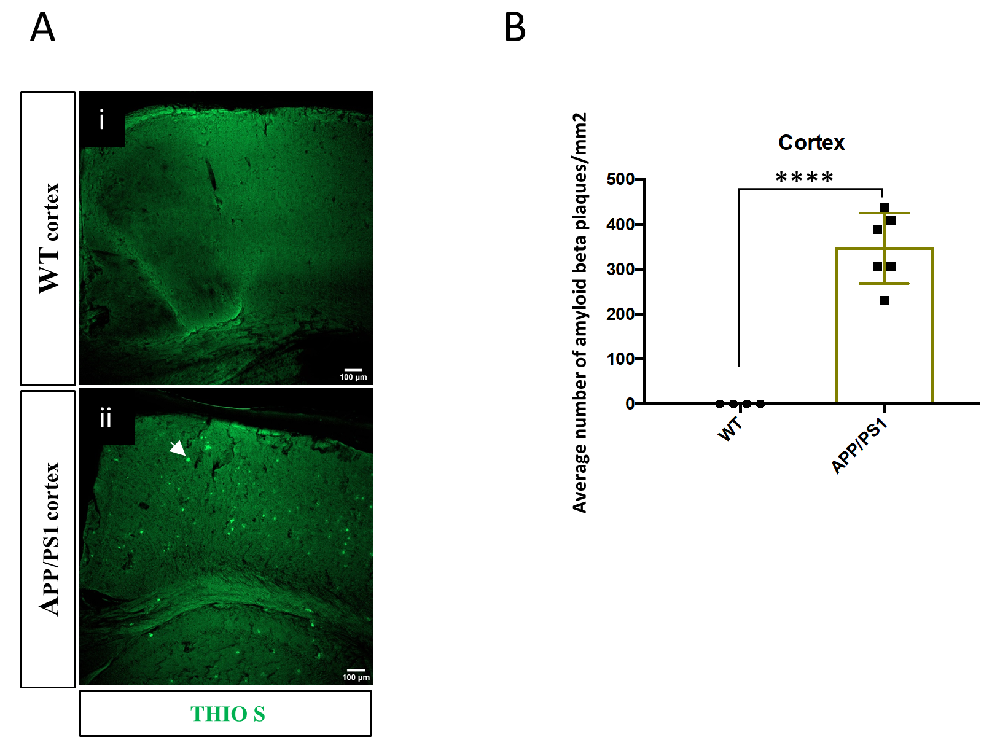
